# Hard-wired visual filters for environment-agnostic object recognition

**DOI:** 10.1101/2024.09.30.615752

**Authors:** Minjun Kang, Seungdae Baek, Se-Bum Paik

## Abstract

Conventional deep neural networks (DNNs) are highly susceptible to variations in input domains, unlike biological brains which effectively adapt to environmental changes. Here, we demonstrate that hard-wired Gabor filters, replicating the structure of receptive fields in the brain’s early visual pathway, facilitate environment-agnostic object recognition without overfitting. Our approach involved fixing the pre-designed Gabor filters in the early layers of DNNs, preventing any alterations during training. Despite the restricted learning flexibility of this model, our networks maintained robust performance even under significant domain shifts, in contrast to conventional DNNs that typically fail in similar conditions. We found that our model effectively clustered identical “classes” across diverse domains, while conventional DNNs tend to cluster images by “domain” in the latent space. We observed that the fixed Gabor filters enabled networks to encode global shape information rather than local texture features, thereby mitigating the risk of overfitting.

**One sentence summary:** Hard-wired Gabor filters enable environment-agnostic object recognition without overfitting.

**Research Highlights:**

- Conventional deep neural networks (DNNs) are vulnerable to input domain variations
- Hard-wired Gabor filters facilitate environment-agnostic object recognition
- Fixed Gabor filters prevent overfitting and facilitate shape-based classifications
- Our model cluster identical “classes” while conventional DNNs cluster by “domain”

## Introduction

Modern deep neural network (DNN) models demonstrate exceptional capabilities in executing various tasks^1,2^ often surpassing human performance^3,4^. However, these models struggle to operate effectively under non-stationary environments^5,6^, particularly when the image domain changes^6–8^. When presented with data from a new distribution (Fig. 1a), DNNs frequently fail to recognize previously learned objects, resulting in a significant loss of performance^5,9,10^ (Fig. 1b, top). This limitation hampers the applicability of DNNs in dynamic scenarios, such as driving in varying weather conditions^11^ or identifying objects under different lighting^12^. To address this issue, several methods have been proposed to enhance the performance of DNNs in dynamic environments. One strategy involves pre-training with extensive datasets^13^ and fine-tuning on new images^14,15^. Another approach leverages upcoming domain information during training to reduce discrepancies between domains^16,17^. However, these data-driven approaches are effective only for a limited number of domains, and the model’s adaptability is significantly challenged by novel datasets unless additional training or data is available. This issue becomes especially critical in dynamic and unpredictable environments.

**Figure 1.**
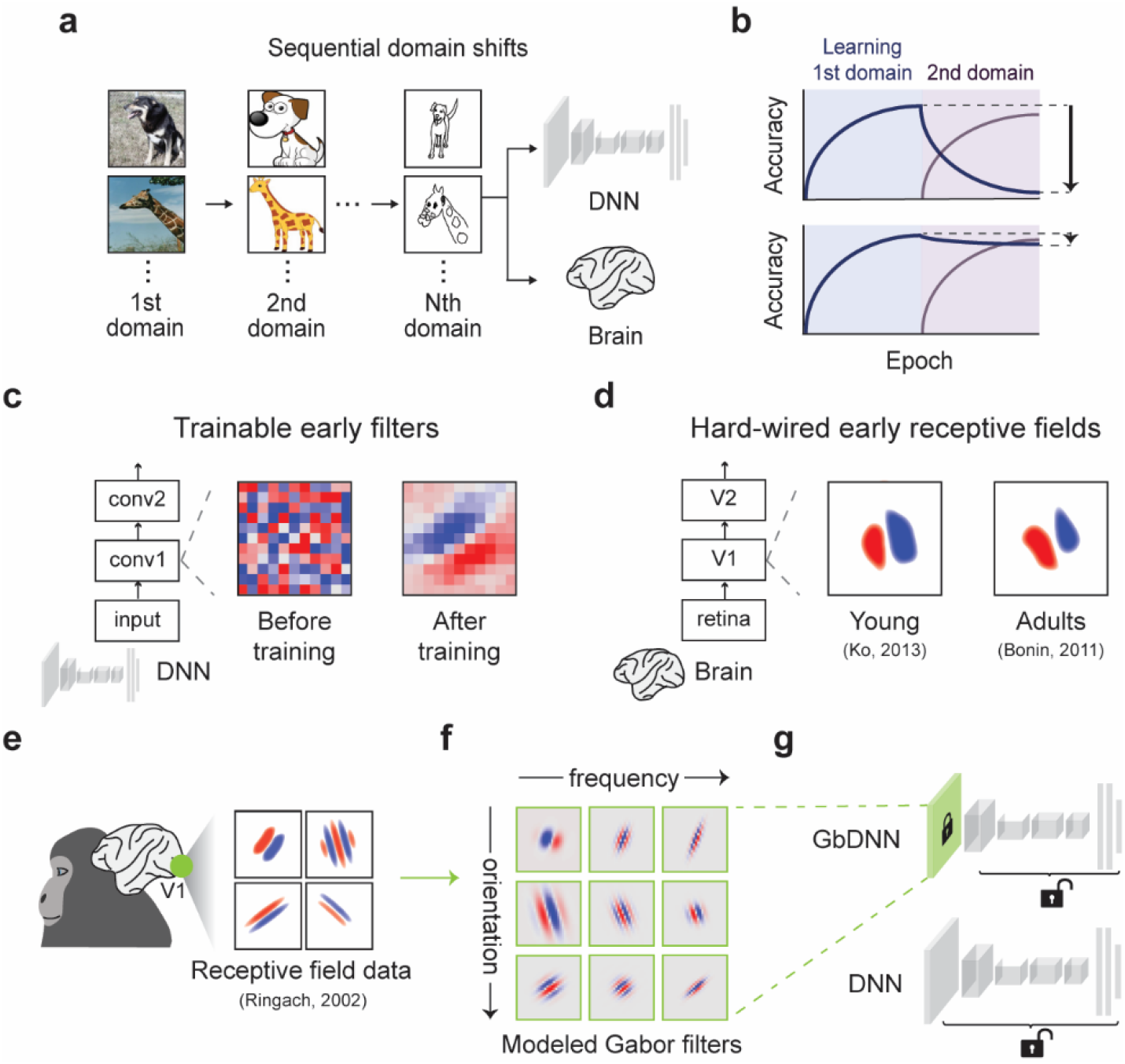
Conventional deep neural networks and brains under dynamic domain shifts. (a) Illustration of learning under sequential domain shifts for conventional deep neural networks (DNNs) and brains. (b) Performance of each network. DNNs exhibit performance degradation (top), whereas brains can retain prior performance (bottom). (c) Early filters in DNNs are randomly initialized and trained depending on the dataset. (d) The brain’s early receptive fields remain stable throughout life. The receptive fields are illustrations inspired by the original papers (Ko, 2013; Bonin, 2011). These are not direct reproductions of the original figures. (e) Illustration of receptive field data. The receptive fields are illustrations inspired by the original papers (Ringach, 2002). These are not direct reproductions of the original figures. (f) Modeled Gabor filters demonstrating variations in orientation and spatial frequency. (g) Our model (GbDNN) contains fixed Gabor filters in the early layer, while conventional DNN is fully trainable.

Given that biological organisms can effortlessly perform diverse tasks across environmental variations^18–20^ (Fig. 1b, bottom), we expected that the structural characteristics of the brain could offer valuable insights. Specifically, we noted that the early layer filters of DNNs often begin as random noise^21,22^ and adapt to match input statistics^23^ (Fig. 1c), whereas the early visual circuitry of the brain—from the retina to the primary visual cortex (V1)—arises spontaneously and maintains its initial structure throughout a lifetime of visual experiences^24–27^ (Fig. 1d). Notably, even before eye-opening, the maturation of this circuitry begins with V1 neurons showing selectivity for specific visual features such as orientations and spatial frequencies^25,28–31^. The functional maps representing these neural tunings, systematically organized within V1^32–36^, remain stable after eye-opening^37–39^ unless optical deprivation occurs^40^. This stability in the early visual pathway may contribute to robust feature encoding and resilience to environmental changes.

Here, we hypothesized that pre-developed filters in the early visual pathway enable environment-agnostic object recognition. To test this, we modeled the receptive fields of V1 neurons (Fig. 1e) as Gabor filters (Fig. S1a) using biologically measured data^41–44^ (Fig. 1f, Fig. S1b) and introduced them into the first layer of the DNN^23^ (Fig. 1g, Fig. S1c). In this model, called GbDNN, the Gabor filters are fixed during training without any modifications. This approach resulted in consistent object recognition across domain shifts, such as from photographs to sketches^45^. We observed that the model clusters object *classes* rather than image *domains* in the latent space, effectively resolving domain discrepancies. We observed that the Gabor filters reduced the dimensionality of representations, preventing overfitting and enabling the networks to utilize global shape information instead of local texture features. Our findings highlight the biological strategies that facilitate reliable visual perception in dynamic environments.

## Results

### Fixed Gabor filters enable environment-agnostic object recognition

To simulate dynamic environments involving domain shifts, we trained networks using input images whose domain changes over time. We utilized the PACS (Photo-Art-Sketch-Cartoon) dataset^45^, which consists of seven common classes distributed across four distinct domains (Fig. 2a, Fig. S2). In this context, “class” refers to the semantic categories such as “dog” and “giraffe”, while “domain” pertains to the style of images, such as “Photo” and “Sketch”. We trained the networks sequentially on one domain followed by another, resulting in twelve unique domain pairs for the continuous training of both DNNs and GbDNNs (Fig. 2b).

**Figure 2.**
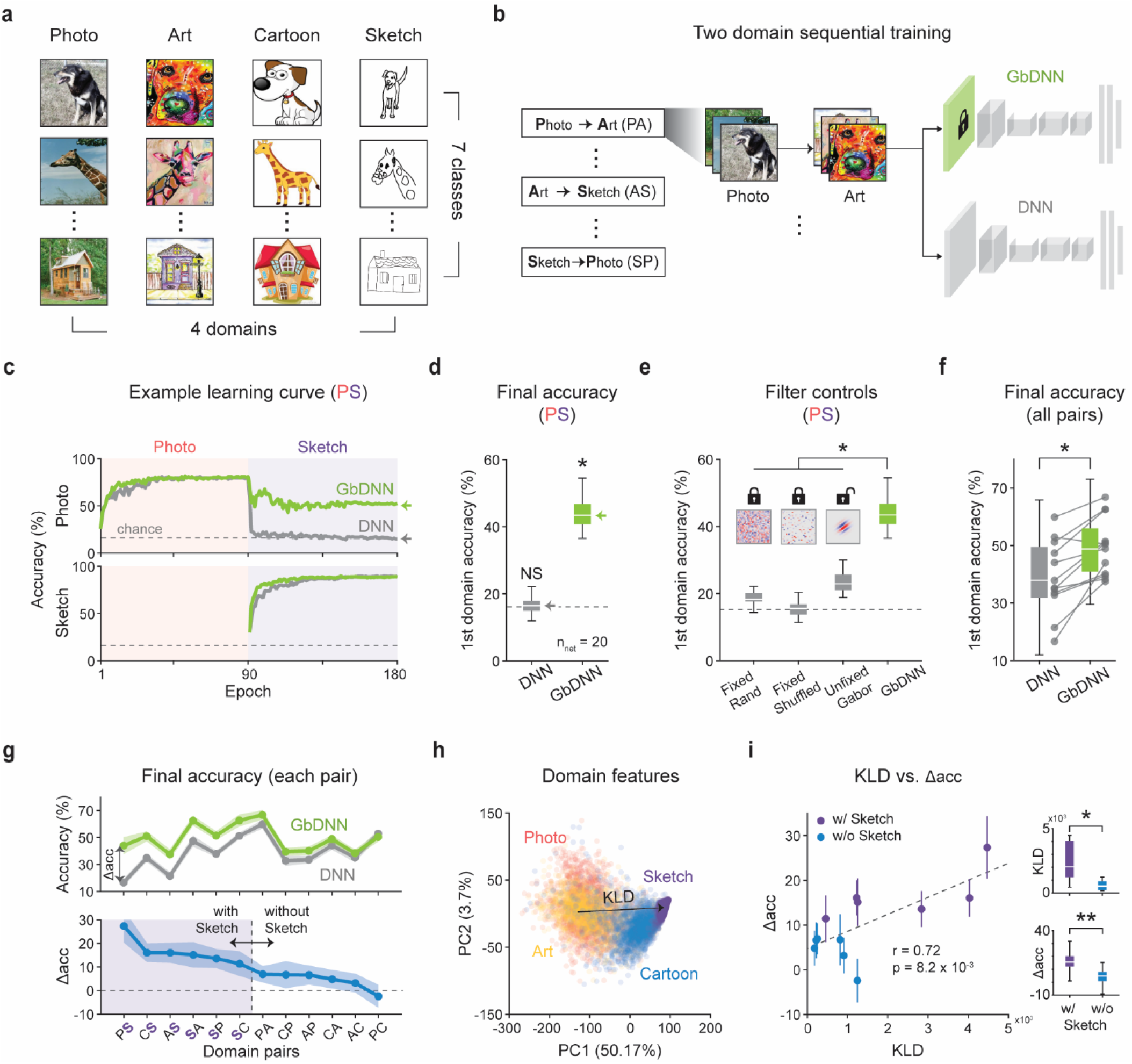
Fixed Gabor filters facilitate environment-agnostic object recognition. (a) Stimuli consisted of four domains and seven common classes^45^. (b) Our task involves sequential training on two different domains from the set. (c) Example learning curve for sequential training on Photo followed by Sketch domains: (top) accuracy for Photo. (Bottom) accuracy for Sketch. (d) Final Photo accuracy after training on Sketch. DNN performance dropped to the chance level (one-sample t-test, n = 20, NS: p = 0.36), while GbDNNs maintained significantly higher performance (paired t-test, n = 20, *p < .001). (e) Final accuracy across different filter controls. GbDNNs outperformed networks with fixed random, fixed shuffled Gabor, and unfixed Gabor filters (one-way ANOVA, n = 20, *p < .001). (f) For all domains, GbDNNs showed higher accuracy in the first domain than DNNs (Wilcoxon signed-rank test, n = 240, *p < .001). The dots represent each domain pair. (g) (top) Final accuracy for each pair. (bottom) Final accuracy differences between GbDNNs and DNNs (Δacc). The larger differences involved the Sketch domain. (h) Domain visualization via PCA, with inter-domain distances measured using Kullback-Leibler divergence (KLD). (i) KLD and Δacc have a significant correlation (Pearson correlation, r = 0.72, p = 8.2 × 10^−3^), with domain pairs involving Sketch showing higher KLD (Wilcoxon rank-sum test, n = 6, *p < .05) and Δacc (Wilcoxon rank-sum test, n = 120, **p < .001). The dots represent each domain pair.

Our results demonstrate that GbDNNs robustly maintained their performance across domain shifts. After training on the first domain, both DNNs and GbDNNs achieved high accuracy optimized for that domain’s dataset (Fig. 2c, top, Fig. S3). This pattern persisted after training on the second domain, with both networks converging on the new dataset (Fig. 2c, bottom). However, training on the second domain led to a notable performance drop for DNNs on the first domain. For instance, DNNs exhibited a complete loss of accuracy on Photo images, which fell to chance level after sequential training on Sketch images (Fig. 2d, DNN vs. Chance, one-sample t-test, n = 20, NS: p = 0.36). In contrast, GbDNNs preserved their initial accuracy significantly above chance level (Fig. 2d, GbDNN vs. Chance, one-sample t-test, n = 20, *p < .001), illustrating the robustness of our model under domain changes.

We found that both the fixed status and Gabor-like structures played crucial roles in the observed effects. To assess the significance of the fixed status of the early layer filters, we simulated networks with unfixed Gabor filters, which were initialized as before, but allowed to be trained. Additionally, we compared networks employing other types of fixed filters, such as randomly initialized or spatially shuffled Gabor filters, to investigate the importance of filter structure. The results showed that GbDNNs significantly outperformed all other conditions in preserving performance on the first domain (Fig. 2e, Unfixed Gabor / Fixed random / Fixed shuffled Gabor / GbDNN, one-way ANOVA, n = 20, *p < .001). These findings indicate that both the fixed status and the Gabor-like structures, which reflect biological characteristics, are essential for maintaining accuracy under domain shifts.

Furthermore, we confirmed that the trend persisted across various domain pairs, with GbDNNs consistently outperforming conventional DNNs (Fig. 2f, DNN vs. GbDNN, Wilcoxon signed-rank test, n = 240, *p < .001). The accuracy differences between GbDNNs and DNNs, Δacc, remained above zero for most domain pairs (Fig. 2g). Notably, Δacc varied depending on the specific domain pairs, being especially pronounced when the Sketch domain was involved (Fig. 2g, bottom). This suggests that larger differences between the trained domains may yield higher Δacc. To further explore this, we measured domain dissimilarity by estimating the Kullback-Leibler divergence^46^ (KLD) using a k-nearest neighbor method^47^ after applying principal component analysis (PCA) to the raw images (Fig. 2h, See Methods for details). Our analysis revealed that domain pairs involving the Sketch domain, which exhibited larger Δacc, also showed higher KLD than the other pairs (Fig. 2i, insets, Wilcoxon rank-sum test, *p < .05, **p < .001). Since the larger domain distances can result in more significant performance degradations, pairs involving Sketch precisely represent the issues we initially aimed to address. By measuring the correlation between KLD and Δacc, we could confirm significant positive correlations between them (Fig. 2i, Pearson correlation, r = 0.72, p = 8.2 × 10^−3^), highlighting the potential strength of GbDNNs under highly varying environments. Overall, GbDNNs demonstrate reliable object recognition under unexpected and abrupt environmental changes, which pose challenges for DNNs.

### Our model spontaneously clustered the same objects across various domains

To understand how GbDNNs maintain performance under domain shifts, we evaluated their ability to produce general object representations across domains. We measured responses from the first fully-connected layer, which is known for encoding categorical information^48^. The t-SNE analysis of the Photo and Sketch pairs revealed that conventional DNNs formed tight clusters based on domain rather than object classes (Fig. 3a, left). This indicates that distinct decision boundaries are required for each domain, limiting their ability to consistently recognize objects in changing environments. In contrast, GbDNNs grouped the same classes together, regardless of domain (Fig. 3a, right), suggesting that they rely on consistent, domain-invariant feature representations.

**Figure 3.**
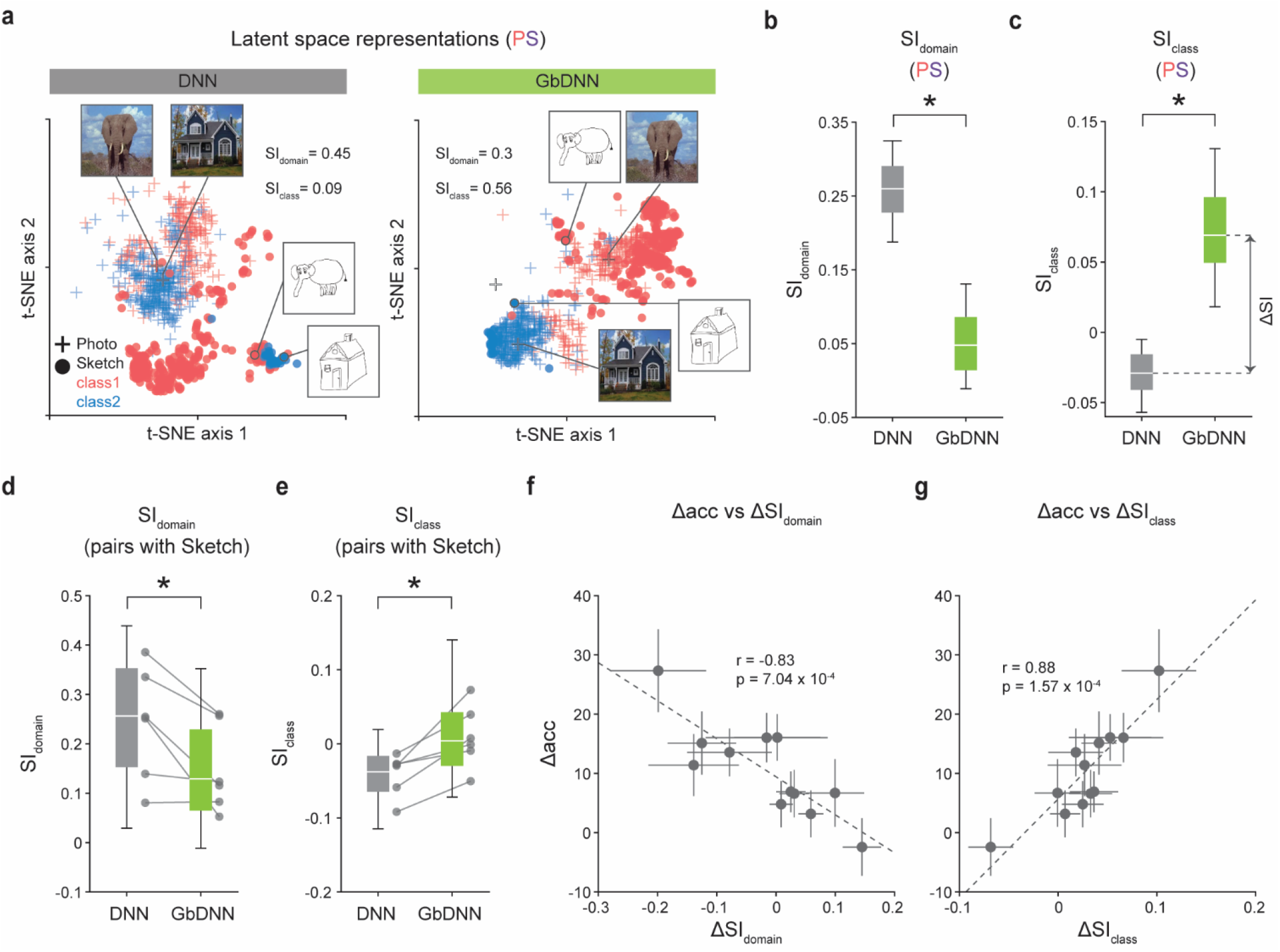
GbDNNs cluster the same classes, while conventional DNNs cluster by domains. (a) Example latent space representations of different classes and domains. DNNs tend to cluster by domain, while GbDNNs cluster by class. (b) DNNs exhibit better domain clustering than GbDNNs, as measured by the silhouette index for domains (SIdomain) (paired t-test, n = 20, *p < .001). (c) GbDNNs showed a higher silhouette index of classes (SIclass) than GbDNNs (paired t-test, n = 20, *p < .001). (d) For domain pairs involving Sketch, DNNs demonstrate higher SIdomain than GbDNNs (Wilcoxon signed-rank test, n = 120, *p < .001). The dots represent each domain pair. (e) For domain pairs involving Sketch, GbDNNs show higher SIclass than DNNs (Wilcoxon signed-rank test, n = 120, *p < .001). The dots represent each domain pair. (f) SIdomain differences and accuracy differences are negatively correlated (Pearson correlation, n = 20, r = -0.83, p = 7.04 × 10^−4^). (g) SIclass differences and accuracy differences showed a significant positive correlation (Pearson correlation, n = 20, r = 0.88, p = 1.57 × 10^−4^).

We quantified this clustering tendency using the silhouette index (SI)^49^, which calculates the ratio of within-class to across-class distances, providing an objective measure across distinct latent spaces^50–52^. Our analysis showed that DNNs exhibited significantly higher SI for domain clusters (SI^domain^) compared to DNNs (Fig. 3b, DNN vs. GbDNN, paired t-test, n = 20, *p < .001), while GbDNNs showed higher SI for class clusters (SI_class_) (Fig. 3c, DNN vs. GbDNN, paired t-test, n = 20, *p < .001). We further explored this trend across domain pairs involving the Sketch domain, previously identified as problematic. While DNNs cluster by domain and achieved a higher SI_domain_ (Fig. 3d, DNN vs. GbDNN, Wilcoxon signed-rank test, n = 120, *p < .001), GbDNNs again demonstrated superior grouping of the same object classes, exhibiting a higher SI_class_ (Fig. 3e, DNN vs. GbDNN, Wilcoxon signed-rank test, n = 120, *p < .001). These findings suggest that GbDNNs more effectively rely on domain-general representations compared to conventional DNNs.

Importantly, the measured SI demonstrated a strong correlation with performance. By analyzing the calculated accuracy differences (Fig. 2g) and the difference between the SI for GbDNNs and DNNs, we found a negative correlation with differences of SI_domain_ (Fig. 3f, Pearson correlation, n = 20, r = -0.83, p = 7.04 × 10^−4^), and positive correlation with differences of SI_class_ (Fig. 3g, Pearson correlation, n = 20, r = 0.88, p = 1.57 × 10^−4^). These strong correlations suggest that the ability to create compact clusters of classes, rather than domains, may serve as a reliable indicator and predictor of the performance differences between GbDNNs and DNNs. Together, these results indicate that hard-wired Gabor filters enable networks to achieve reliable perception by generating universal object representations that are independent of image domains.

Next, to examine the direct role of Gabor filters in processing multiple domains, we analyzed the geometry of the representations in the first layer. We measured the activations of the first convolutional layer from networks trained on the same images (Fig. S4a). Given that the responses are three-dimensional, we flattened them and applied PCA to extract eigenvalues (Fig. S4b, left). A large eigenvalue indicates that the corresponding axis explains a substantial amount of variance in the representations. Thus, a sharper drop in the eigenvalue spectrum suggests that the representation tends to lie in low-dimensional subspaces, requiring fewer axes to explain the total variance (Fig. S4b, right). We found that fixed Gabor filters reduce the dimensionality of representations; GbDNNs exhibited a steeper drop in eigenvalues (Fig. S4c) compared to conventional DNNs. To quantify this, we calculated the quadratic entropy of these spectra, which measures the effective dimension^53,54^ of the representations (See Methods for details). The results showed that GbDNNs consistently displayed lower effective dimensions than conventional DNNs across various domains (Fig. S4c, DNN vs. GbDNN, n = 20, paired t-test, *p < .001). By integrating data from all domains, we confirmed that GbDNNs exhibit a steep drop in the spectra and lower effective dimensions (Fig. S4d, DNN vs. GbDNN, n = 80, Wilcoxon signed-rank test, *p < .001). This suggests that GbDNNs inherently produce low-dimensional representations, which may reduce the risk of overfitting to the training images.

### Fixed Gabor filters enable shape-biased object classifications

Given that GbDNNs produce general representations, an important question arises: what specific features do GbDNNs encode? One hypothesis is that GbDNNs encode global shapes that remain invariant across domain changes, akin to biological organisms that prioritize shape recognition^26,55^, in contrast to conventional DNNs which focus more on local textures^56,57^. To test this hypothesis, we generated images that capture shape information while removing local textures (Fig. 4a, Fig. S5) using Frangi filters^58^, which detect edges through hessian-based methods (See Methods for details). Additionally, we created texture-preserving images by randomly shuffling local patches, retaining local information while disrupting global shapes^59^. We then presented these image sets to networks trained on the original images to assess their object classification bias.

**Figure 4.**
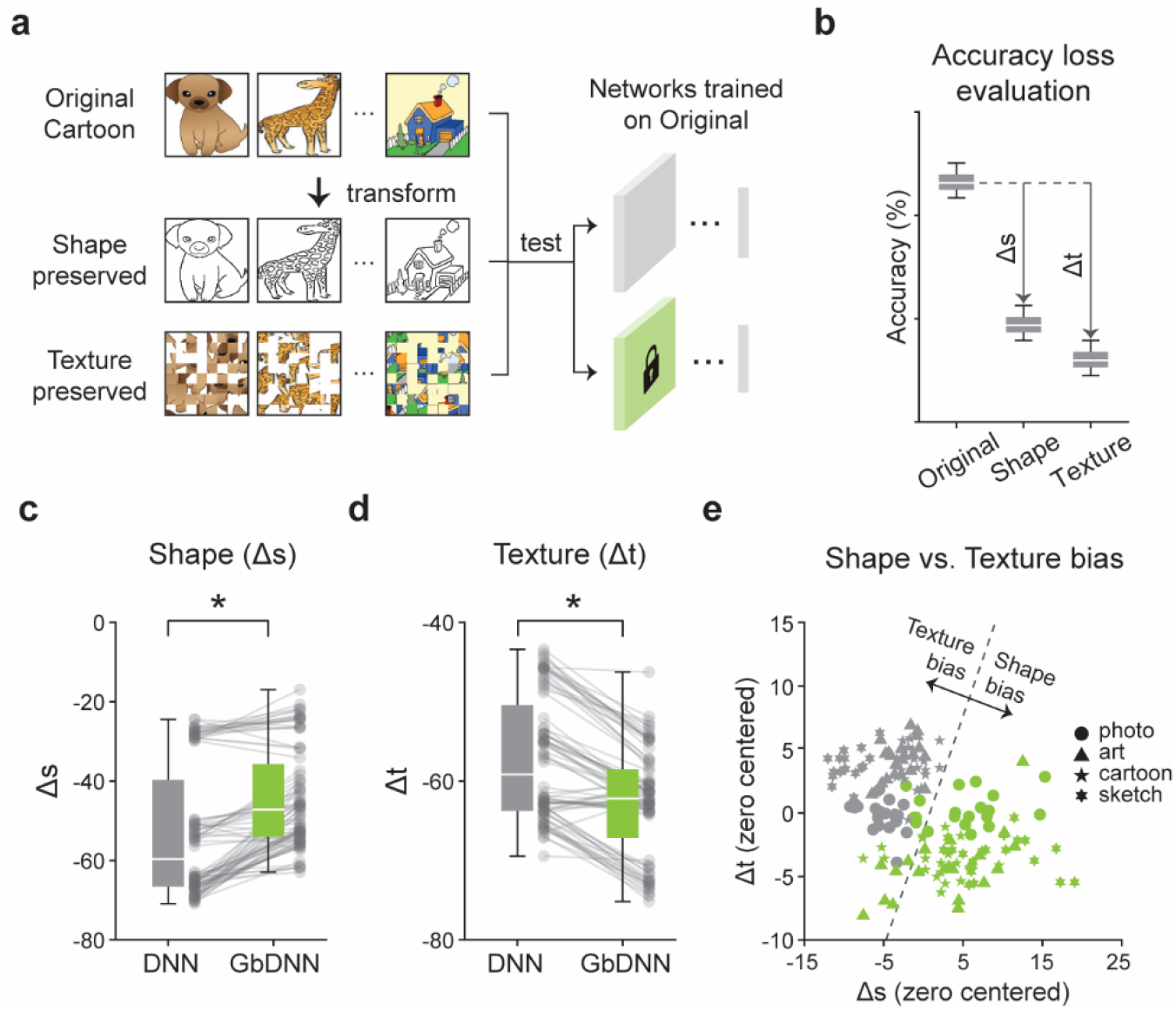
GbDNNs show shape-biased object recognition. (a) Illustration of testing shape- and texture-biased object recognition. Networks trained on original images were evaluated on shape images, created using Frangi filters, and texture images, created by shuffling local patches. (b) Performance loss was measured for shape (Δs) and texture (Δt) images. (c) For shape images, GbDNNs show lower accuracy loss than DNNs (Wilcoxon signed-rank text, n = 80, *p < .001). The dots represent each network. (d) For texture images, DNNs show lower accuracy loss than GbDNNs (Wilcoxon signed-rank text, n = 80, *p < .001). The dots represent each network. (e) Scatter plots of zero-centered Δs and Δt demonstrate that GbDNNs are biased towards shape-based object classification, whereas DNNs exhibit a texture bias.

Our results demonstrated that GbDNNs exhibit shape-biased object recognition. We measured the accuracy loss of shape and texture images from the accuracy for original images (Fig. 4b) and compared their tendencies. As a result, GbDNNs experienced a smaller drop in accuracy for shape images across all domains compared to DNNs (Fig. 4c, DNN vs. GbDNN, paired t-test, n = 80, *p < .001), indicating that they better encode shapes and maintain performance. Conversely, for texture images, DNNs exhibited a smaller accuracy drop than GbDNNs (Fig. 4d, DNN vs. GbDNN, paired t-test, n = 80, *p < .001), suggesting that DNNs prioritize texture information. When plotting both shape and texture accuracy loss together, we confirmed that GbDNNs exhibit a stronger bias toward shapes, while DNNs show a greater bias toward textures (Fig. 4e). These findings emphasize our model’s capability to encode global and domain-general information effectively.

## Discussion

Our model proposes a biologically plausible framework for integrating physical circuits into networks. Given that the receptive fields of the retina and lateral geniculate nucleus (LGN) resemble Gaussian filters^60^, local convergence projections can facilitate the spontaneous emergence of Gabor filters in the primary visual cortex (V1) through the superposition of ON- and OFF-centered Gaussian receptive fields^31,61^. Furthermore, it has been suggested that functional maps, such as those representing orientation or spatial frequency, can arise from the interference of regularly spaced retinal mosaics, even in the complete absence of learning^34–36,62^. These findings lend support to the incorporation of a fixed Gabor filter bank as a viable strategy for brain-like object recognition. Our results indicate that GbDNNs exhibit robustness to domain shifts by generating domain-general representations. Such features are frequently observed in neurophysiological studies^63,64^, where invariant activations occur in response to face images presented in various styles^63^. Additionally, GbDNNs exhibit shape-biased object classification, a phenomenon observed in both adults^56,65^ and infants^66,67^ data. In contrast to other approaches that attempt to implement shape bias into networks through image augmentations^68^ or adversarial training^56^, our approach proposes a biologically plausible mechanism for shape-biased perception without further training or post-processing.

Our method demonstrates advantages in real environments that often encounter unexpected, substantial changes. Specifically, GbDNNs inherently reduce domain discrepancies and mitigate the risk of overfitting. The clustering results (Fig. 3a, left) reveal distinct separations between the Photo and Sketch domains, represented by crosses and filled circles, respectively, indicating that conventional DNNs map different domains into separate subspaces. In contrast, GbDNNs display overlapping clusters (Fig. 3a, right), which facilitate the maintenance of decision boundaries and further reduce overfitting. Moreover, our method is particularly effective under extreme domain shifts (Fig. 2i). While the performance difference between GbDNNs and DNNs is minimal in cases where domains are similar (e.g., Art to Cartoon), GbDNNs outperform DNNs when domains vary significantly (e.g., Photo to Sketch). In such scenarios, the incorporation of hard-wired Gabor filters can preserve network performance.

What factors contribute to the functional advantages of our model? We propose that several physical characteristics of hard-wired Gabor filters provide insights. First, Gabor filters function as spatial band-pass filters, effectively reducing redundant high-frequency content and allowing the network to concentrate on global features. The use of multiple Gabor filters, each tuned to different orientations and spatial frequencies, enables the capture of structural information within images, ensuring that each filter identifies distinct aspects of the local image region without redundancy. Furthermore, the presence of fixed filters in the early layers helps stabilize the distribution of features in subsequent layers. By maintaining consistent initial feature extraction, the network can achieve more stable intermediate representations, thereby mitigating the effects of covariate shifts throughout the network.

Another significant advantage of GbDNNs is their data training-free nature, enabling universal application across various image domains. First, GbDNNs are not susceptible to dataset biases. Traditional approaches such as transfer learning^15^, often involve pre-training on labeled datasets such as ImageNet^13^. Despite extensive training on large image collections, transfer learning models frequently inherit biases from the pre-trained data, exemplified by texture bias in ImageNet-trained networks^56,57^. In contrast, our model effectively addresses these issues through a brain-inspired, data-free methodology. Moreover, GbDNNs operate without the need for future information. Conventional domain adaptation strategies^7,17,69,70^ aim to train a network on both a “source” domain and a “target” domain. This typically requires access to “target” domain images during training to align representations, which is often impractical for real-world applications where future images are unknown. By eliminating reliance on training images, our model can operate effectively in diverse environments in a data training-free manner.

Our method provides a practical advantage due to its straightforward implementation, even within physical networks. Specifically, the diverse range of Gabor filters can be easily initiated using Gaussian filters. Previous studies have suggested that Gabor filters with diverse orientations and spatial frequencies can emerge spontaneously from the interference of regularly spaced Gaussian filters^34–36,62^. In the context of physical networks, such as neuromorphic computing^71^, this can be achieved by hard-wiring Gaussian filters at the input stage and arranging them in a regular pattern, allowing simple interference to generate the complete set of Gabor filters. Additionally, our method does not require extra computations or resources. Unlike recent machine learning approaches that necessitate additional storage ^72,73^ or supplementary neural networks ^69^, our model minimizes resource requirements by fixing the early layers, reducing the number of parameters. This simplicity and efficiency render GbDNNs well-suited for applications in physical neural networks and other devices where memory and computational resources are constrained.

In summary, our findings suggest a brain-inspired strategy, utilizing inherent kernels from the early visual pathway, can facilitate environment-agnostic visual perception. These results may provide insights into achieving reliable object recognition for both biological and artificial neural network systems.

## Methods

### Receptive fields

The illustrations of receptive fields presented in Fig. 1c (left), 1c(right), 1d are created based on the descriptions and data provided by Ko et al. (2013), Bonin et al. (2011), Ringach (2002), respectively. These are not direct reproductions of the original figures. Full credit for the original data and inspiration for these illustrations goes to the original authors.

### Gabor filters

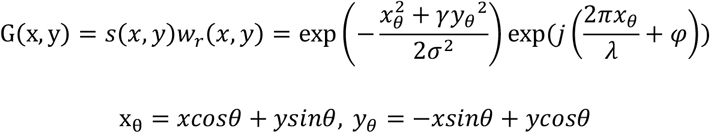

The Gabor filter is defined by a complex sinusoidal wave localized in both frequency and orientation. It can be expressed as the multiplication of a complex sinusoid, known as the carrier (*s*(*x, y*)), and the Gaussian function as the envelope (*w*_*r*_ (*x, y*)). First, *σ* determines the entire size of the filter and affects the spatial frequency. Second, *γ* controls the ratio between the x-axis and y-axis dispersion. Third, *θ* defines the specific orientation tuning. Fourth, *φ* represents the phase value. Finally, *λ* determines the wavelength of the filter.

In the first convolutional layers of GbDNNs, 96 types of Gabor filters with distinct parameters were used (Fig. S1a). In order to mirror the biological characteristics, spatial frequency^44^, orientation^43^, and Gaussian envelope^41^ were randomly sampled from macaque V1 data distributions (Fig. S1b). This ensures each network has a unique set of Gabor filters while maintaining consistent distributional statistics (Fig. S1c). As the spatial frequency was denoted as cycles per degree, we set one visual degree to correspond to 15 pixels to ensure the stable performance of models. Additionally, filter sizes were resized from 11 × 11 pixels to 30 × 30 pixels to avoid aliasing and capture Gabor filter properties more effectively.

### Neural network model

We used AlexNet^23^ as a biologically inspired neural network model to simulate the visual system. AlexNet consists of five convolutional layers with rectified linear units (ReLU), a pooling layer for feature extraction, and three fully-connected layers for classification. Inputs of 227 pixels in width and height and seven readouts were utilized for the simulation. To enable comparison with GbDNNs, the filters of the first convolutional layer were also modified to 30 × 30 pixels in conventional DNNs. The network was initialized with all bias values set to zero, weight values randomly sampled from a Gaussian distribution with a zero mean, and the standard deviation set to balance the strength of input signals across the layers^22^. To train networks with various image domains, we employed training hyperparameters that yielded the best training results: optimizer—adaptive moment estimation (Adam), number of epochs = 90 (per each domain), batch size = 64, momentum = 0.9, gradient decay factor = 0.9, squared gradient decay factor = 0.999, initial learning rate = 0.0001, learning rate decay factor = 0.1 for every 30 epochs. Other parameters followed values from the original paper, such as stride for 4 pixels, zero-padding, and dropout properties. To enhance robustness against object positions, all images were randomly flipped along the x-axis and translated within 10% of their original size in both the x and y axes. All simulations were conducted on twenty networks with different initializations.

### Image dataset

We utilized the PACS (Photo-Art-Cartoon-Sketch) dataset^45^ to simulate dynamic environments that require continuous adaptation. This dataset comprises four domains, each containing seven common classes: dog, elephant, giraffe, guitar, horse, house, and person. For each domain, 1670, 2048, 2344, and 3928 images were in photo, art, cartoon, and sketch, respectively (Fig. S2). Among them, we used 90% of the images for training and the remaining images for validation. The accuracies reported in the main figures represent validation accuracy. For sequential training tasks, two domains were used for training out of the four available domains, resulting in 12 unique combinations. Each domain was trained for 90 epochs, resulting in a total of 180 epochs. The final accuracy for the first domain, as shown in Figure 2, was calculated at the 180^th^ epoch after the full training process was completed.

### Domain distance calculation

To measure domain distances, we employed Kullback-Leibler divergence (KLD)^46^ which quantifies the divergence between two probability distributions. The formula for KLD is given by:

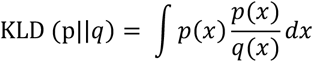

where *p*(*x*) and *q*(*x*) are probability density functions for two continuous densities p and p. Since our images are represented as samples rather than distributions, we estimated the probability density of distributions using the k-nearest neighbor method proposed by Kozachenko-Leonenko^47^. Suppose p and *q* are continuous densities, and let {*X*_1_, …, *X*_*n*_} and {*Y*_1_, …, *Y*_*nn*_} be i.i.d. samples drawn independently from p and *q*, respectively. The estimation of KLD becomes:

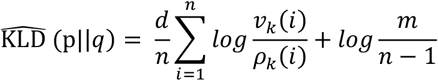

where n and m are number of samples in two densities p and *q*, respectively, and *ρ*_*k*_ (*i*) be the Euclidean distance between *X*_*i*_ and its k-nearest neighbor in {*X*_*j*_}_*j*≠*i*_ and *v*_*k*_ (*i*) be k-nearest neighbor in {*Y*_*i*_}. The hyperparameter k was set to 10 considering the high dimensionality of image PC spaces. All analyses were conducted after PCA was conducted on the original images, and we used axes that explained 99% of the variance.

### Clustering evaluation

To evaluate learned representations, we measured the responses from the first fully connected layer after the entire training sequence. For example, in networks trained on Photo followed by Sketch domain, we fed both Photo and Sketch images together to the trained networks and obtained activations. Then, the dimension reduction was performed along the feature axis using t-SNE (Fig. 3a). The scatter plots in Fig. 3a represent data from a single network with two example classes (elephant and house), while the silhouette index (SI) results (Fig. 3b-e) were calculated from twenty networks to provide the summary statistics.

The silhouette index (SI) was used to evaluate the quality of clustering. Basically, it estimates the consistency of data clustering by considering both the within-cluster distance and the across-cluster distance for each point. The silhouette index for the *i-th* point (*SI*_*i*_) can be expressed as

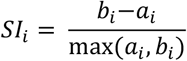

where *a*_*i*_ and *b*_*i*_ refer to intra-class and inter-class distance, respectively. The intra-class distance is defined as the average distance with points from the same cluster, and the inter-class distance is defined as the average distance with points from the other clusters. For domain and class SI calculation, cluster labels were assigned based on domain labels and class labels, respectively.

### Shape and texture image generation

We utilized Frangi filter^58^, which is commonly used for vessel enhancement, to generate the shape images. Frangi filter enhances tubular structures within 2D images by calculating eigenvalues of the Hessian matrix. The eigenvalues are combined into a single measure to enhance contours within images. We chose the Frangi filter for two reasons: (1) it helps eliminate the effect of low-level features like scale or position, and (2) it includes a hyperparameter for shape thickness, which suited our simulations better than traditional edge detectors that often produce very thin, one-pixel-wide edges. For implementation, we used the function “fibermetric” from MATLAB to apply this filter. Object polarity was set as dark and line width as 5. The pixel values that show smaller than 60% intensity were set to zero in order to reduce noisy lines for robust generations.

To generate texture images, we shuffled local patches after diving into the original images. Specifically, we first set the width of each local patch whose width and height are 28 pixels (Fig. S4a), yielding 16 patches for a 227 × 227 pixel image. The patches were shuffled randomly and then joined together to create the shuffled images. To avoid incorrect accuracy from the class imbalance, we used the same number of images for both shape and texture images.

### Effective dimension

In order to evaluate the generalizability of representations, we calculated the effective dimension (ED)^53^ of the covariance matrix using the following formula:

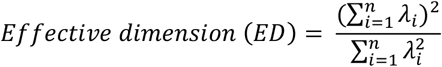

where *λ*_*i*_ refers to the *i-th* eigenvalue. First, the raw activations were calculated by feeding images into the first convolutional layer. Each activation was vectorized since conv1 generates three-dimensional activations. PCA was then conducted on the activation matrix. The obtained eigenvalues were sorted in decreasing order and scaled them by dividing with the largest eigenvalue for visualization (Fig. S4). Finally, the quadratic entropy was computed from the raw eigenvalues to determine the effective dimension of the matrix.

### Statistics

All sample sizes, significance levels, and statistical methods are indicated in the corresponding text and figure legends. The paired t-test was used for all analyses comparing 20 DNNs and 20 GbDNNs, except for comparing with the chance level (one-sample t-test, Fig. 2d), comparing with multiple networks (one-way ANOVA, Fig. 2e), comparing for all domain pairs (Wilcoxon signed-rank test, Fig. 2f, 3d, e, 4c, d; Wilcoxon rank-sum test, Fig. 2i). All box plots in Fig. 2d-f, i, 3b-e, 4c, d, S4 c, d indicate the inter-quartile range (IQR between Q1 and Q3) of the dataset; the horizontal line reveals the median and the whiskers correspond to the rest of the distribution (Q1 − 1.5*IQR, Q3 + 1.5*IQR). The error bars and shaded areas in Fig. 2g, i, 3f, g, S4c indicate the standard deviation across 20 networks. The correlations were measured by Pearson correlation in Fig. 2i, 3f, g.

## Acknowledgments

This work was supported by the National Research Foundation of Korea (NRF) grants (NRF-2022R1A2C3008991 to S.P.) and by the Singularity Professor Research Project of KAIST (to S.P.).

## Author contributions

S.P. conceived of the project. M.K., S.B., and S.P. designed the model. M.K. performed the simulations. M.K., S.B., and S.P. analyzed the data. M.K. drafted the manuscript and designed the figures. S.P. wrote the final version of the manuscript. All authors discussed and commented on the manuscript.

## Supplementary Figures

**Figure S1.**
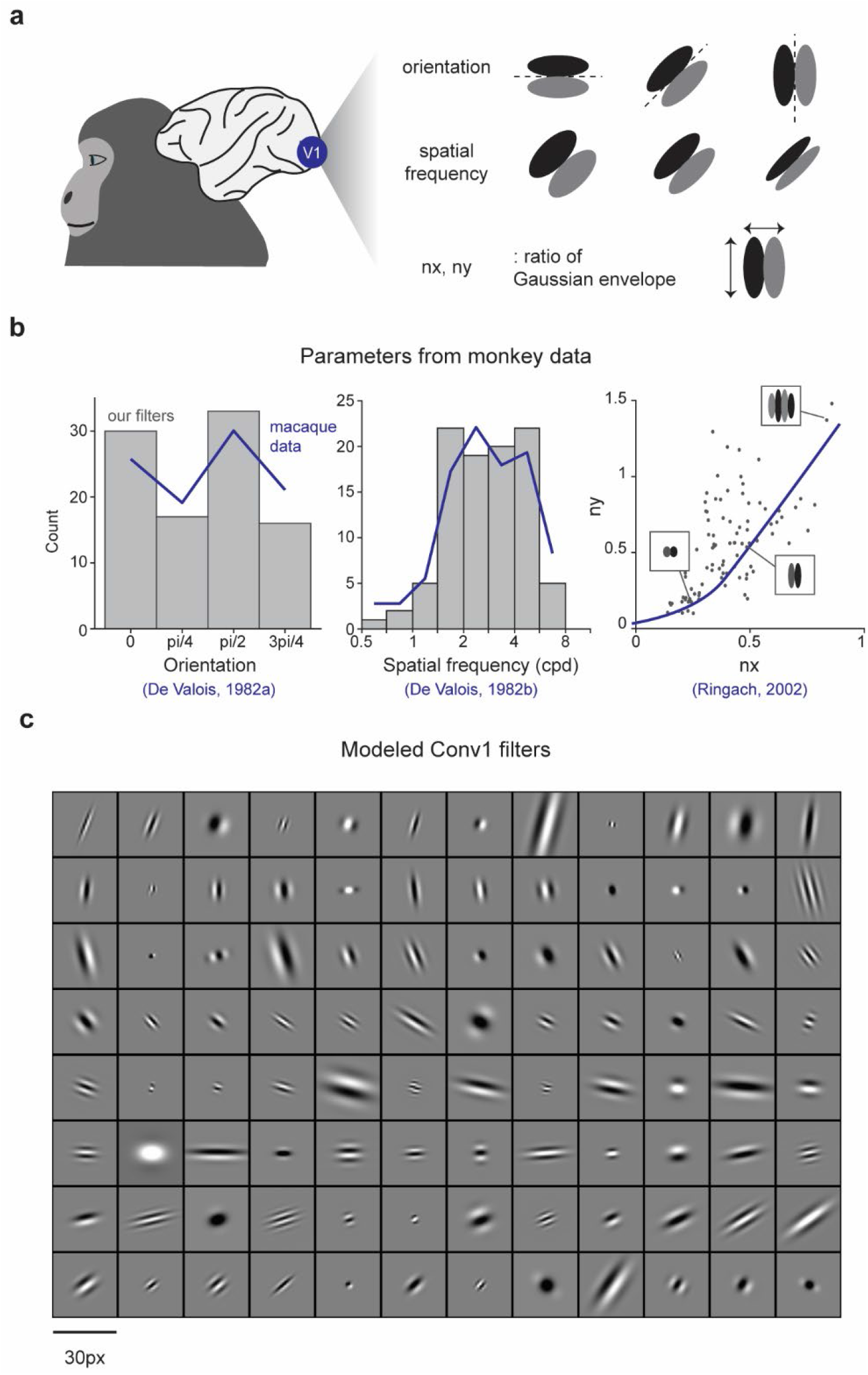
Overview of filter statistics. (a) Illustration of Gabor filter parameters, including orientation, spatial frequency, and the normalized Gaussian envelope. (b) The parameters of our filters were sampled from macaque data^41,43,44^. (c) Example conv1 filters that were hard-wired into the GbDNN.

**Figure S2.**
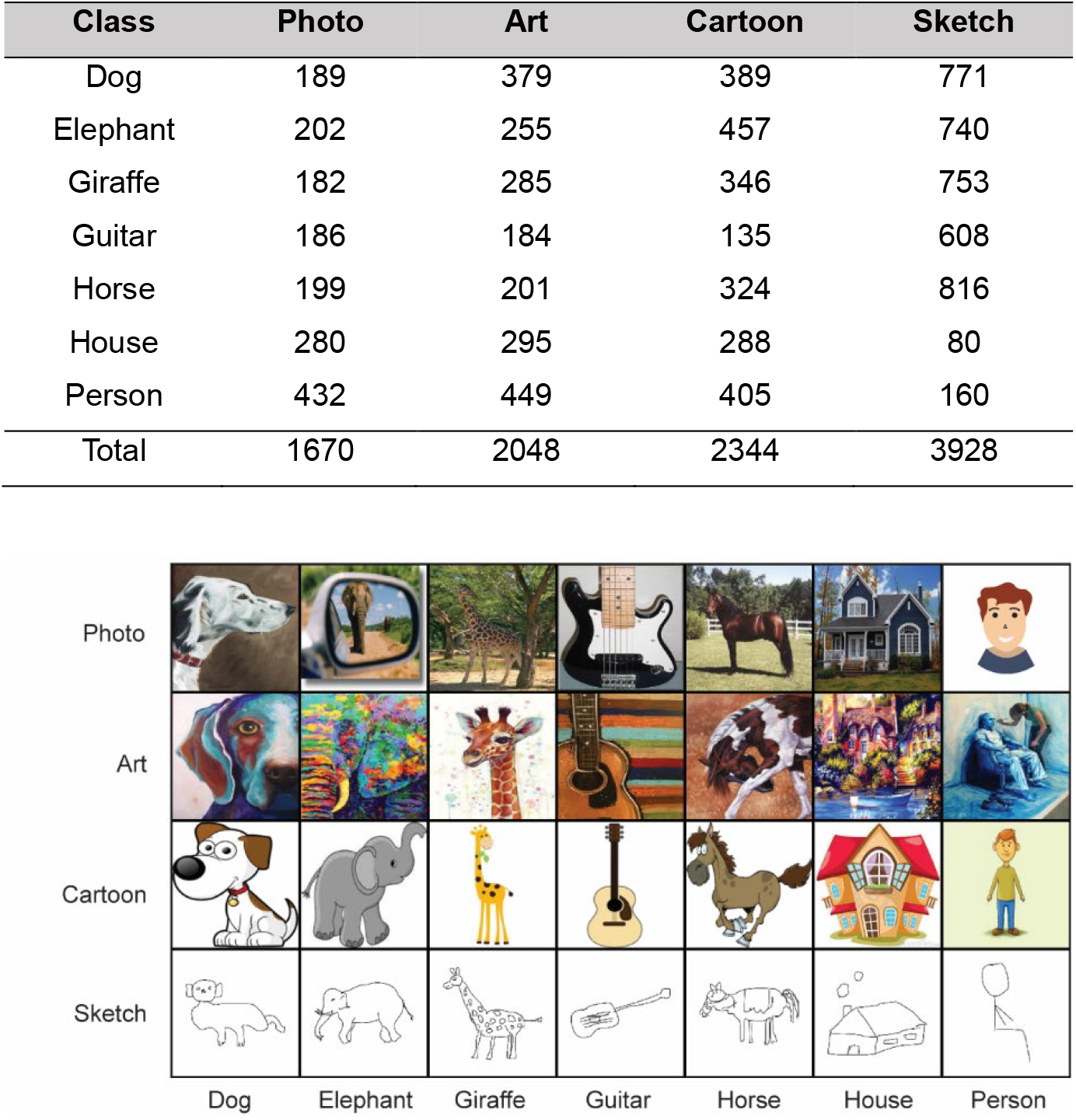
Overview of the stimulus set. Example images from each domain and class of the PACS dataset^45^. All domains share the same object categories. The “Person” class in the “Photo” domains was represented by an illustration instead of an original photo, in compliance with bioRxiv policy.

**Figure S3.**
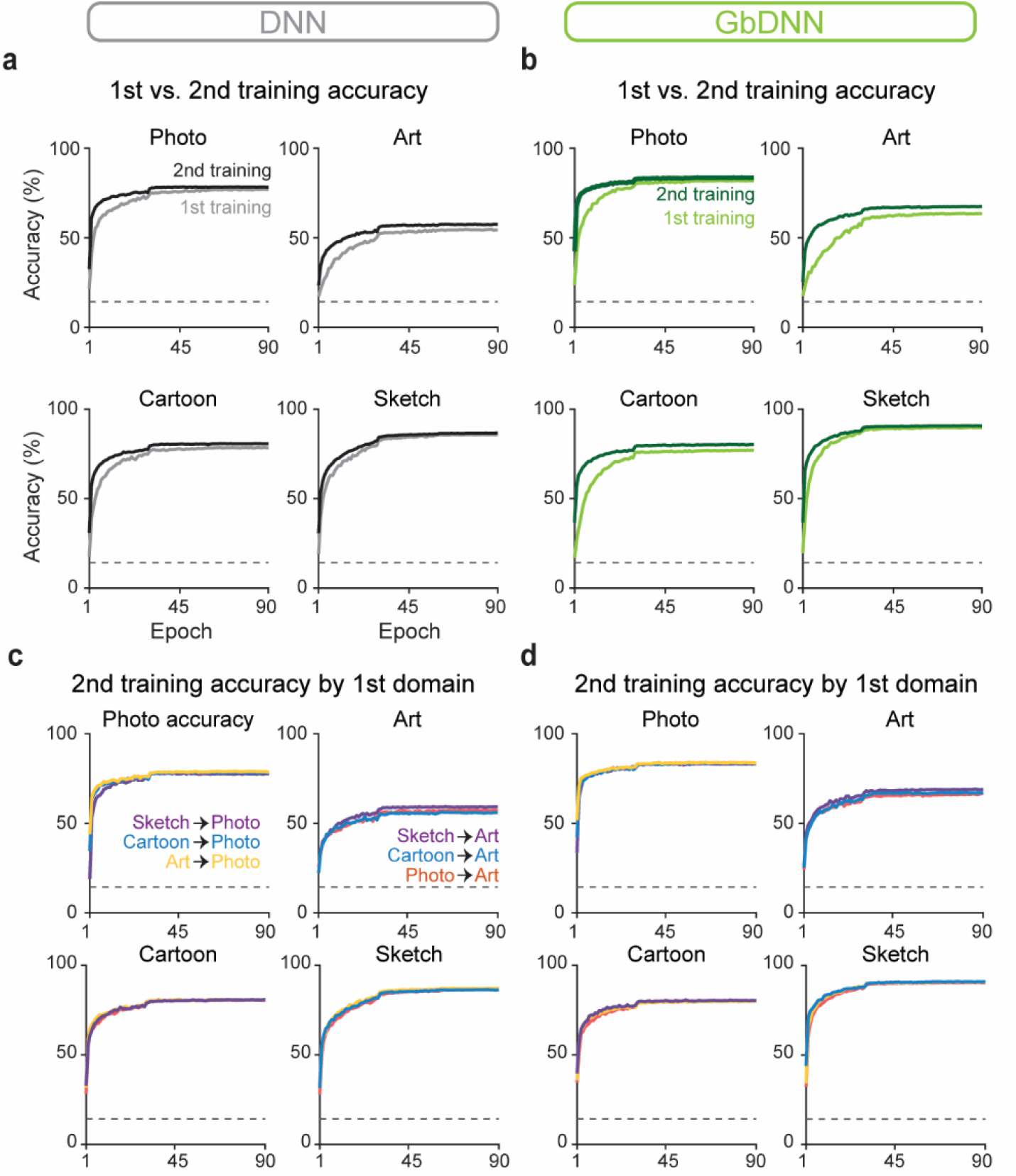
Convergence of training across orders and domains. (a-b) Training accuracy for each domain at different training phases. All networks were converged to comparable accuracies across different domains, regardless of the training phases. (c-d). Second-phase training accuracy for each domain by different types of first-trained domains. The second-phase accuracies showed similar convergence, regardless of the initial domain trained.

**Figure S4.**
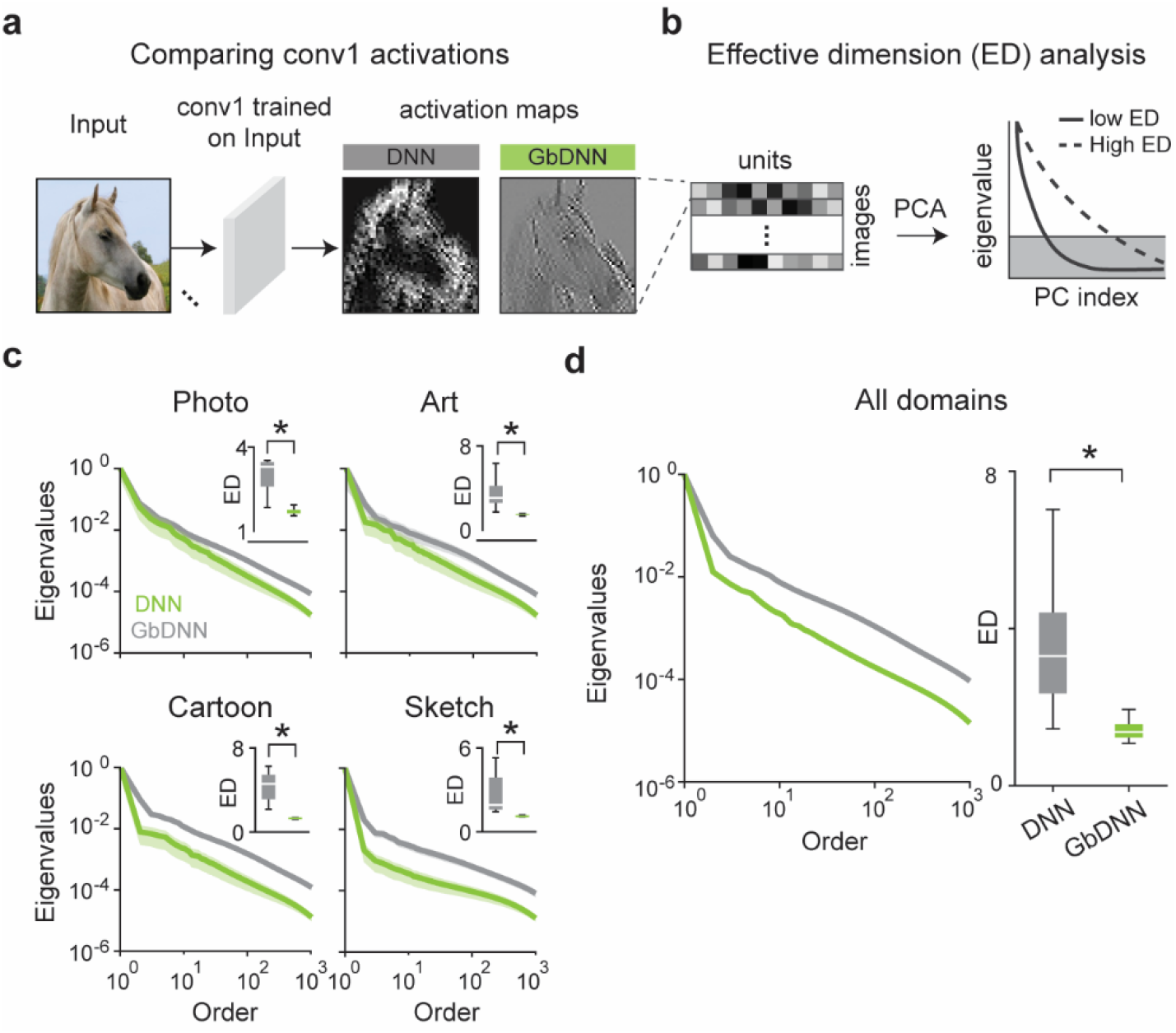
Fixed Gabor filters reduce the dimensionality of representations. (a) Comparison of conv1 activations between DNN and GbDNN. (b) Illustration of the effective dimension analysis. After conducting PCA on conv1 activations, eigenvalues were sorted in descending order. A sharper drop of eigenvalues indicates a lower effective dimension. (c) GbDNNs exhibited a sharper eigenvalue drop and lower effective dimension than DNNs, regardless of the training domain (paired t-test, n = 20, *p < .001). (d) Across all domains, GbDNNs show a shaper drop of eigenvalues and lower effective dimension than DNNs (Wilcoxon signed-rank test, n = 80, *p < .001).

**Figure S5.**
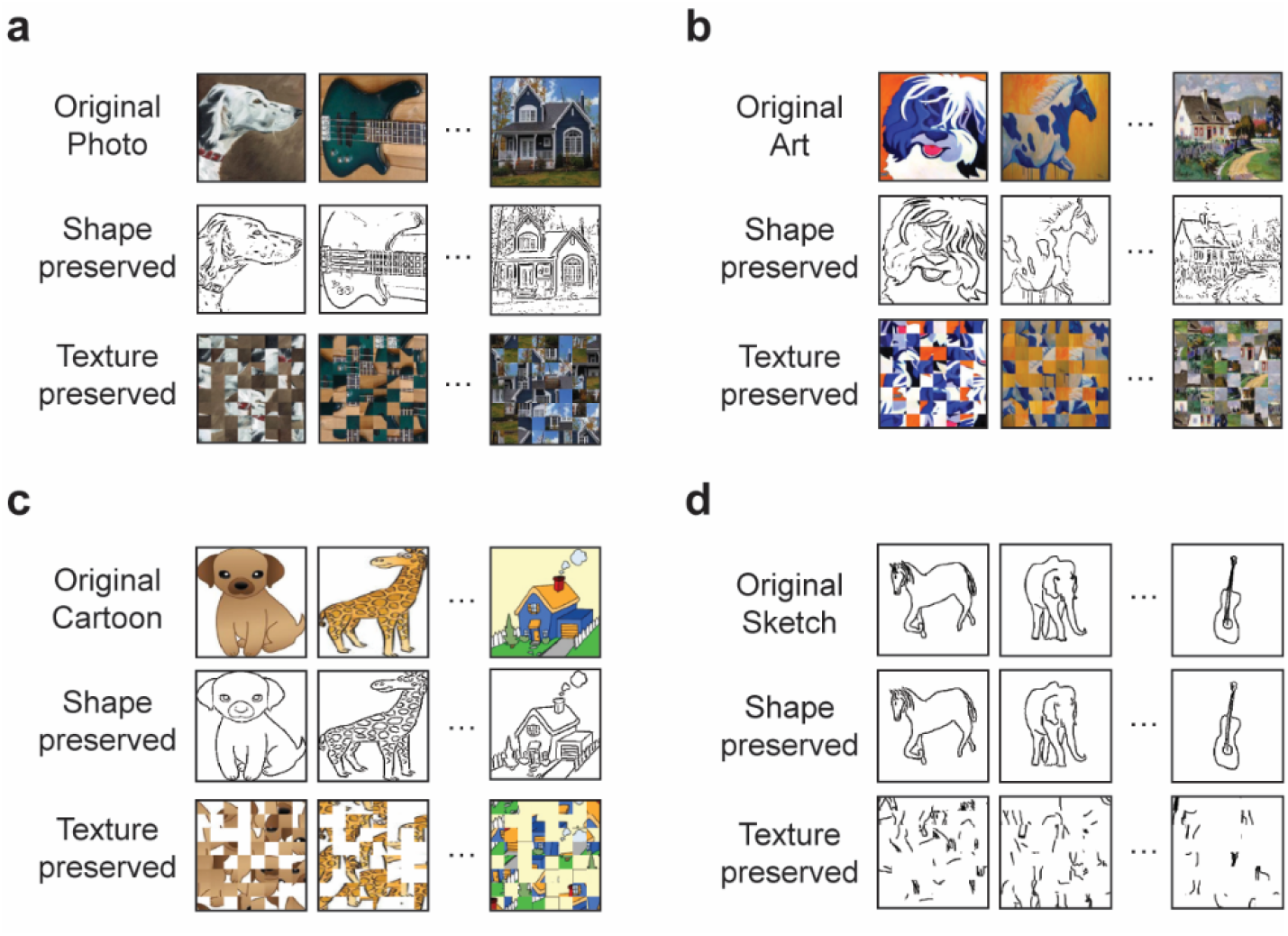
Texture and shape image examples for each domain. (a-d) Example shape and texture images obtained from the original Photo, Art, and Cartoon images. Shape images were generated using the Frangi filter, and texture images were created by shuffling local patches from the original images.

